# Impaired Signaling of NF-κB and NRF2 in CX3CR1-Deficient Microglia: Implications in Tauopathies

**DOI:** 10.1101/346304

**Authors:** Sara Castro-Sánchez, Ángel J. García-Yagüe, Sebastian Kügler, Isabel Lastres-Becker

## Abstract

TAU protein aggregation is the main characteristic of neurodegenerative diseases known as tauopathies. Low-grade chronic inflammation is also another hallmark that indicates crosstalk between damaged neurons and glial cells. We have demonstrated that neurons overexpressing TAU^P301L^ release CX3CL1, which activates anti-inflammatory NRF2 signalling in microglial cells *in vitro* and *in vivo.* However, the potential role of CX3CR1 in the context of tauopathies and its implication in neuroinflammation are poorly described. In this work we show that CX3CL1 activates the pro-inflammatory pathway as an early response mediated by the transcription factor NF-κB through the activation of mitogen-and stress-activated protein kinase-1 (MSK-1). At a functional level, CX3CR1-deficient microglia show decreased expression of NRF2, impaired cell migration and deficiency of phagocytosis. The relevance of these findings is evident in a tauopathy model, where the treatment with an inducer of NRF2, sulforaphane, is able to modulate astrogliosis but not microgliosis. These findings suggest that CX3CR1/NRF2 axis is essential in microglial activation associated with tauopathies and that polymorphisms have to be taken into account to development of therapeutic strategies

## INTRODUCTION

Neurodegenerative diseases such as Alzheimer’s disease (AD), frontotemporal dementia with parkinsonism linked to chromosome 17 (FTDP-17), progressive supranuclear palsy, Pick’s disease and corticobasal degeneration are characterized by the deposition of microtubule-associated protein TAU and known collectively as tauopathies. These diseases share important clinical, pathological, biochemical and genetic characteristics, although the molecular events that lead from conformation changes in normal TAU protein to neuronal dysfunction and cell death are essentially unknown and are probably diverse (Williams, 2006). Besides neuronal degeneration, it is also known that the neuron environment contributes to this event, where glial cells play a crucial role (Ransohoff, 2016). In this context, neuroinflammation with a reactive morphology of astrocytes and microglia, together with low to moderate levels of proinflammatory markers is a key factor in tauopathies. In this regards, the implication of inflammation in neurodegeneration could suggest non-cell autonomous processes. This has been confirmed by several evidences: first, neurodegenerative diseases have genetic hallmarks of autoinflammatory disease (Richards, Robertson et al., 2018). Second, very recently it has been described that immune memory in the brain is an important modifier of neuropathology (Wendeln, Degenhardt et al., 2018). Third, the discovery of genetic variants only expressed by microglia among the central nervous system (CNS) like TREM2, CD33 and CR1, that have been associated with AD, FTLD and other neurodegenerative diseases (HollingworthHarold et al., 2011, Lambert, Heath et al., 2009, Li & Zhang, 2018, Simsvan der Lee et al., 2017, Villegas-Llerena, Phillips et al., 2016, Wes, Sayed et al., 2016). Among these genes, polymorphisms in CX3CR1 influence disease progression but not risk in Alzheimer’s disease as well as in amyotrophic lateral sclerosis, two diseases characterized by neuroinflammation (Lopez-Lopez, Gamez et al., 2014, Lopez-Lopez, Gelpi et al., 2017). CX3CR1 is the receptor for the chemokine fractalkine (CX3CL1) and is a critical signaling pathway for microglia-neuron crosstalk (Lauro, Catalano et al., 2015, Mecca, Giambanco et al., 2018). Interestingly, the CX3CL1/CX3CR1 axis is implicated in the regulation of cognitive functions and synaptic plasticity, particularly in the hippocampus. Disruption in this pathway has been associated with impaired neurogenesis and in aged rats there are decreased levels of hippocampal CX3CL1 protein (Bachstetter, Morganti et al., 2011, de Miranda, Zhang et al., 2017). These data indicate that the CX3CL1/CX3CR1 axis declines with age, an essential key factor for neurodegeneration. As to tauopathies, we previously described that hippocampal neurons expressing human TAU^P301L^ mutant protein produce CX3CL1 and the hippocampus of patients with AD also exhibited increased expression of CX3CL1 in TAU-injured neurons that recruit microglia (Lastres-Becker, Innamorato et al., 2014). Therefore, it is very important to understanding the involvement of CX3CR1 in chronic neuroinflammation and cognitive impairment. Related to neurodegenerative diseases, it has been reported that in some models CX3CR1-deficiency appears to be neuroprotective (Cho, Sun et al., 2011, Lastres-Becker et al., 2014, Maphis, Xu et al., 2015) but in others CX3CR1-deficiency protects against amyloid β-induced neurotoxicity (Dworzak, Renvoise et al., 2015, Fuhrmann, Bittner et al., 2010, Lee, Xu et al., 2014), indicating the need for more exhaustive studies of the molecular mechanisms involved in the signalling of the CX3CL1/CX3CR1 axis and its implications in tauopathies.

In this context, one of the mechanisms triggered by CX3CL1 is the activation of the transcription factor Nuclear Factor (erythroid-derived 2)-like 2 (herein referred as NRF2). NRF2 recognizes an enhancer sequence termed antioxidant response element (ARE) that is present in the regulatory regions of over 250 genes (ARE genes) such as antioxidant enzymes and biotransformation reactions, lipid and iron catabolism and mitochondrial bioenergetics (Dinkova-Kostova, Kostov et al., 2018). Furthermore, evidences showed that NRF2 is also implicated in the modulation of inflammatory processes through crosstalk with the transcription factor NF-κB, the principal regulator of inflammation (Cuadrado, Martin-Moldes et al., 2014). Additionally, it has been described that NRF2 is essential in proteostasis, which modulates the proteasome and autophagy processes (Lastres-Becker, Ulusoy et al., 2012, Pajares, Jimenez-Moreno et al., 2016). Therefore, modulation of NRF2 activity has the potential to alter neurodegenerative disease course (Burnside & Hardingham, 2017, Cuadrado, Kugler et al., 2018, Lastres-Becker, 2017, Lastres-Becker, Garcia-Yague et al., 2016). Interestingly, previous work showed that NRF2-and CX3CR1-knockout mice did not express heme oxygenase 1 (HO1) in microglia, which led to increased microgliosis and astrogliosis in response to neuronal TAU^P301L^ expression, showing many similarities between both genotypes related to tauopathy (Lastres-Becker et al., 2014).

Therefore, in this work we analysed in depth the molecular mechanisms implicated in the CX3CL1/CX3CR1 axis in inflammation and also the consequences for tauopathies.

For this purpose, we analysed the involvement of the CX3CL1/CX3CR1 axis in the neuroinflammatory process and determined the molecular pathways implicated. Next, we evaluated the role of the CX3CR1 receptor in the modulation of NRF2 signature and its relevance in phagocytosis. Finally, to evaluate the role of CX3CR1 in neurodegeneration, we determined whether the treatment with sulforaphane, a NRF2 activator, could modulate neuroinflammation in a tauopathy mouse model in absence of CX3CR1, which would indicate the relevance of CX3CR1 and NRF2 loss of function polymorphisms in developing therapeutic strategies for humans.

## RESULTS

### CX3CL1 induces NF-κB-p65 and pro-inflammatory cytokines expression in microglial cells

In order to assess whether CX3CL1 was able to activate pro-inflammatory signalling pathways in microglial cells, we first analysed the subcellular distribution of the transcription factor NF-κB-p65, a master regulator of the inflammatory response. Immortalized microglial cells (IMG) were maintained under serum-free conditions for 16 h and then stimulated with CX3CL1 (100 nM) and data were collected at different time points. After 30 min of treatment, p65 starts to translocate to the nucleus (Figure 1A) and reaches a maximum at 120 min after treatment, indicating that CX3CL1 is able to activate pro-inflammatory response as an early event. This was corroborated by the observation that mRNA and protein levels of the pro-inflammatory factors IL-1β, IL-6 and TNF-α increased after CX3CL1 treatment (Figure 1C-D). Our results indicate that CX3CL1 is able to activate the pro-inflammatory pathways as an early molecular event in microglial cells.

**Figure 1:**
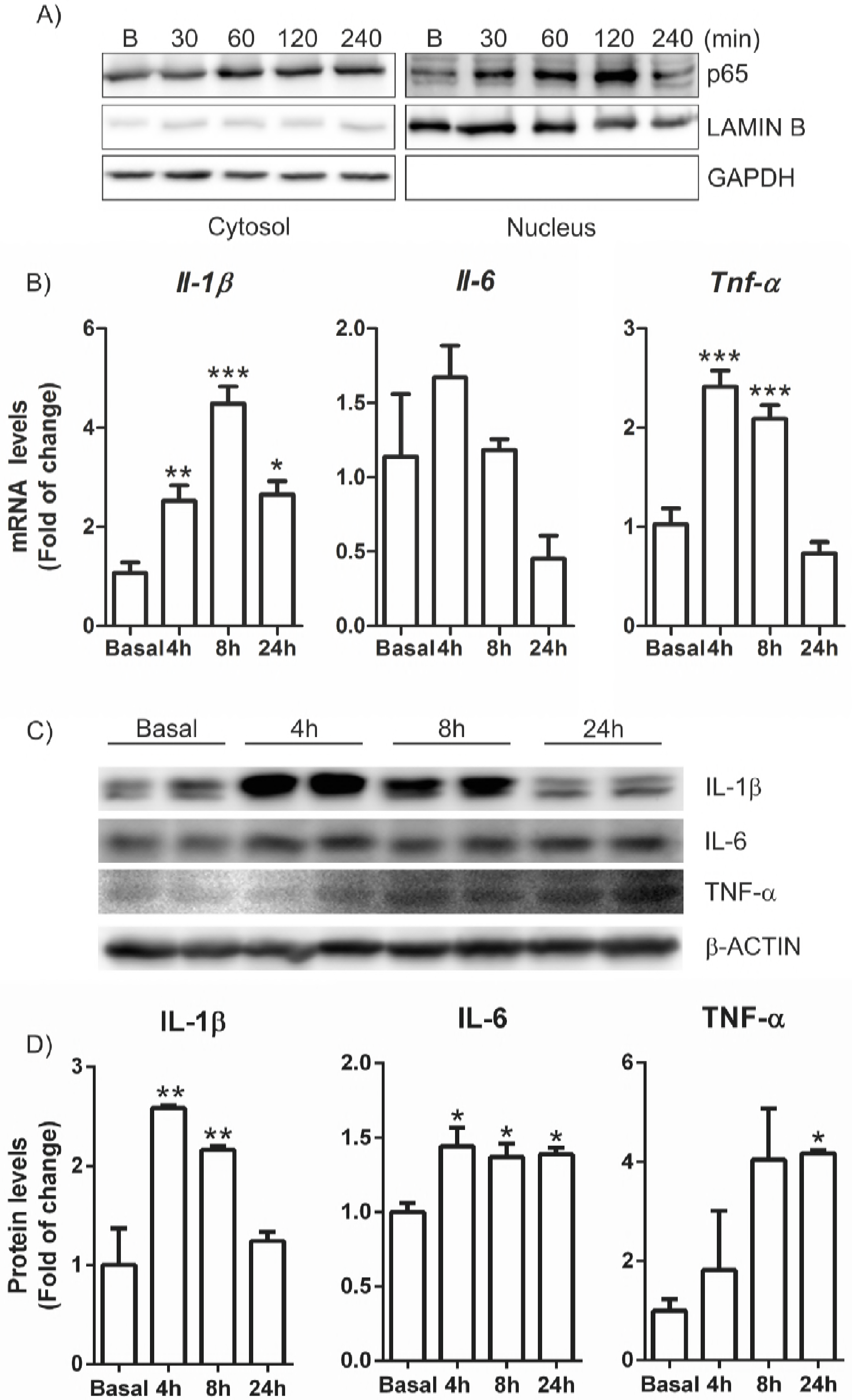
CX3CL1 induces NF-κB signalling in IMG microglia. Cells were incubated in the presence of recombinant CX3CL1 (100 nM) for 30, 60, 120 and 240 min as indicated. (**A**) Analysis by subcellular fractionation in immunoblots: NF-κB levels (top); LAMIN B level was used as nuclear protein loading control (middle); GAPDH levels used as cytosol protein loading control (bottom). (**B**) Cells were incubated in the presence of recombinant CX3CL1 (100 nM) for 4, 8 and 24 h and quantitative real-time PCR determination of messenger RNA for *Il-1β, Il-6* and *Tnf-α* was analysed and normalised by *Actb* (β-Actin) messenger RNA levels. (**C**) Cells were incubated in the presence of CX3CL1 for 4, 8 and 24 h. Immunoblot analysis in whole cell lysates of protein levels of IL-1β, IL-6, TNF-α and β-actin as loading control. (**D**) Densitometric quantification of representative blots from C normalized for β-ACTIN levels. Bars indicate mean of n=3-4 samples ± SEM. Asterisks denote significant differences *p<0.05, **p<0.01 and ***p < 0.001, comparing the indicated groups with the basal condition according to a one-way ANOVA followed by Newman-Keuls post-test.

### CX3CL1 triggers several kinase pathways that converge to MSK1-dependent NF-κB signalling activation

Although the implication of several signalling pathways involved in the modulation of CX3CL1 actions has been described (Al-Aoukaty, Rolstad et al., 1998, Sheridan & Murphy, 2013), such as PI3K/AKT (Lastres-Becker et al., 2014, Lyons, Lynch et al., 2009), most of the data were obtained in neuronal cell types (Sheridan & Murphy, 2013). Therefore, we investigated what signalling pathways could be activated by CX3CL1 when we used a kinase proteome profiler assay (Figure 2A) in immortalized microglia (IMG) cells treated with CX3CL1 (100 nM) for 1 h. Interestingly, the results indicate that CX3CL1 treatment induced the activation of three members of MAPK: extracellular signal-regulated kinase (ERK), p38, and c-Jun N-terminal kinase (JNK) (Figure 2B). The MAPKs are crucial players in cell signalling. MAPKs transmit a broad range of extracellular signals to mediate various intracellular responses (Popiolek-Barczyk & Mika, 2016). We reported previously that CX3CL1 was able to induce the PI3K/AKT/GSK-3β pathway (Lastres-Becker et al., 2014) in BV2 cells, which modulates the transcription factor NRF2 activation. These results were corroborated in IMG cells (Fig. 2B) and in a time-dependent experiment (Suppl Fig.1). Interestingly, our results show that CX3CL1 also activated 5′-prime-AMP-activated protein kinase (AMPK), which has critical roles in regulating growth and reprogramming metabolism, as well as autophagy and cell polarity (Mihaylova & Shaw, 2011). Furthermore, it has been reported that microglial polarization to the M2 phenotype and reduction of oxidative stress are mediated through activation of AMPK and NRF2 pathways (Wang, Huang et al., 2018), what confirms our results. CX3CL1 also induced epidermal growth factor-receptor (EGF-R), which is involved in microglial migration (Qu, Liu et al., 2015). As to mammalian target of rapamycin (mTOR), CX3CL1 stimulates its phosphorylation at Ser 2448, which is implicated in the binding to both, raptor and rictor (Rosner, Siegel et al., 2010). To date, there is no much evidence of mTOR implication in microglial mechanisms, although mTOR selectively controls microglial activation in response to pro-inflammatory cytokines and appears to play a crucial role in microglial viability (Dello Russo, Lisi et al., 2009). Our data shows that CX3CL1 treatment activates SRC-family tyrosine kinases like Yes, Fgr, Lck and Lyn in IMG. Perhaps, this activation is associated with the acquisition of such pro-inflammatory phenotype of the microglia (Socodato, Portugal et al., 2015). All these data indicate that the CX3CL1/CX3CR1 pathway is involved in different signalling processes such as cell polarization, migration and activation of cytokines.

**Figure 2:**
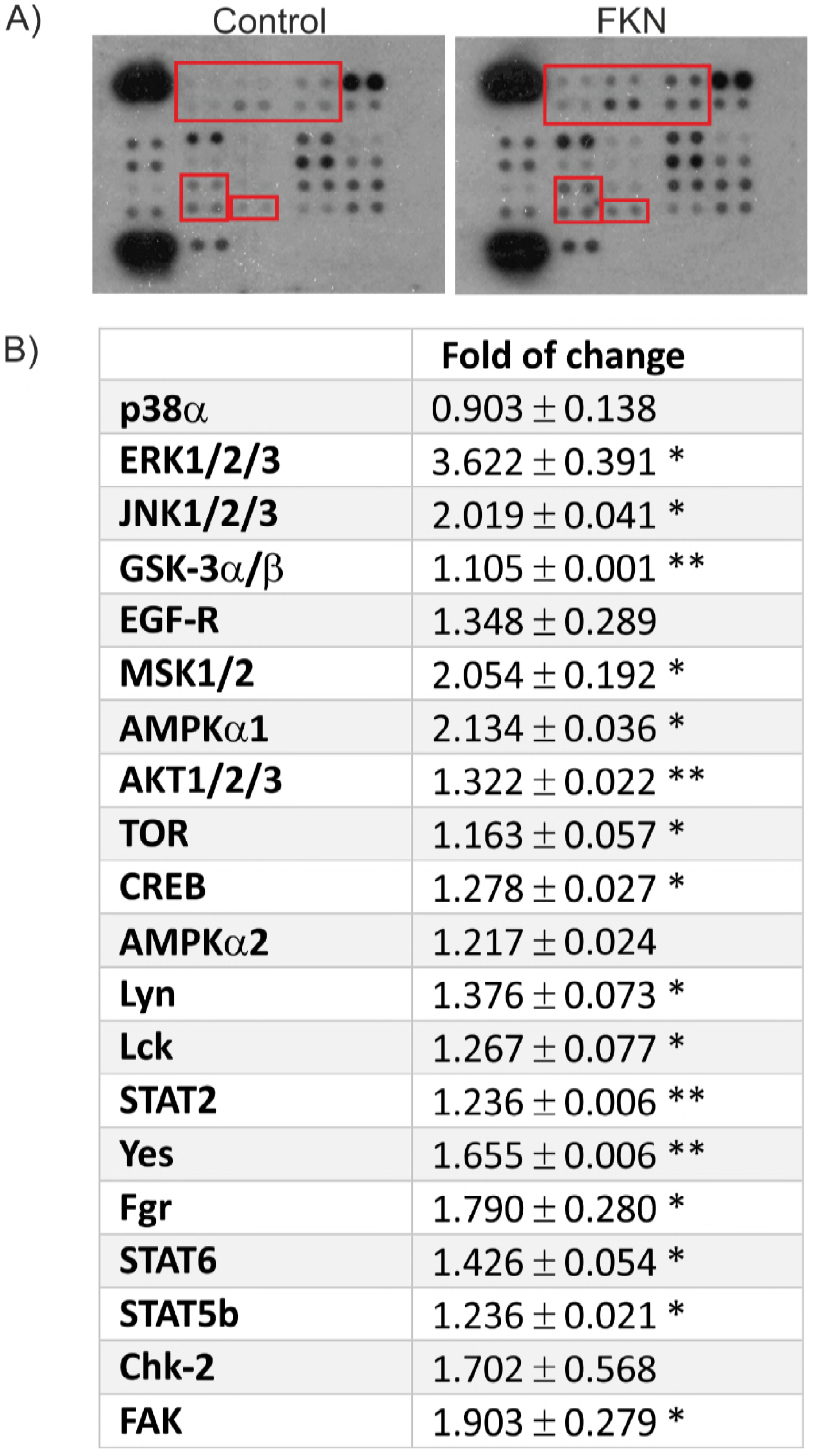
Protein phosphorylation profiling induced by CX3CL1 in primary and immortalized microglial cells. Cells were treated with CX3CL1 (100 nM) for 1h. (**A**) Array spots were visualized in accordance with the manufacturer’s instructions. The intensity of each spot was measured as described in “Materials and Methods”. Red squares indicate similarities between primary and immortalized microglial cells. (**B**) The table shows the relative fold change of proteins with significant difference upon CX3CL1 treatment, setting 1 for control (no treatment of CX3CL1). The data are shown as an average of two individual sets of sample.

It is remarkably, that ERK1/2 and p38-MAPK signalling pathways are involved in NF-κB transactivation (Kefaloyianni, Gaitanaki et al., 2006) through mitogen-and stress activated kinase-1 (MSK1). Our results show that CX3CL1 activated MSK1 (Fig.2B) in both cell types. Hence, we investigated whether MSK1 downregulation could modulate the pro-inflammatory mechanisms triggered by CX3CL1 stimulation in microglia. For this purpose, we knocked down MSK1 with short interfering RNAs (siRNAs). IMG cells were transfected with siRNA against MSK1 (Fig. 3A) or scramble siRNA as control. Partial knockdown of the protein (about 50%, Fig. 3A) was enough to abolish the induction of mRNA levels of *Il-1β, Tnf-α* and *iNos* produced by stimulation with CX3CL1 (Fig. 3B-D, respectively), indicating the relevance of MSK1 activation in the pro-inflammatory signalling pathway induced by CX3CL1.

**Figure 3:**
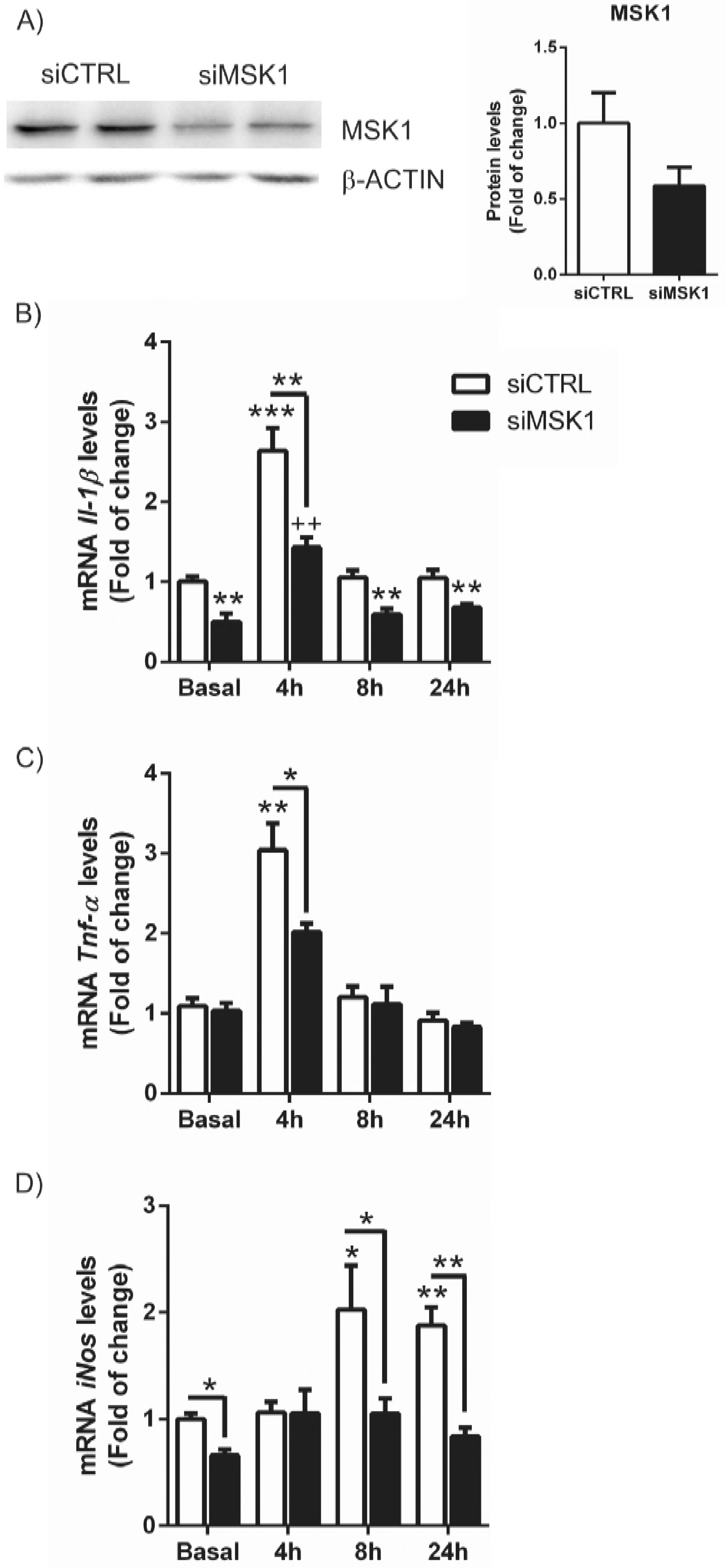
CX3CL1 activates NF-κB signalling in a MSK-1-dependent manner. (**A**) IMG cells were transfected with siRNAs for MSK-1 or with a control scrambled siRNA. Cells were lysed at 48 h after siRNA transfection. Whole-protein lysates were immunoblotted with specific antibodies as indicated in the panels. (**B-D**) mRNA levels of *Il-1β, Il-6* and *Tnf-α*, respectively, were analysed after siRNA knockdown and CX3CL1 (100 nM) treatment for 4, 8 and 24h. Values are mean±SEM (n=4). Statistical analyses were performed with one-way ANOVA followed by Newman-Keuls multiple-comparison test. *p<0.05, **p<0.01, and ***p<0.001 versus IMG cells transfected with scrambled siRNA.

### CX3CR1-deficient primary microglial cells present impaired levels of the transcription factors NF-κB and NRF2 signalling

Our data demonstrate that CX3CL1 activates NF-κB signalling pathway in microglial cells and the implication of MSK1 in this process. Moreover, MSK1 downregulation modulates the basal mRNA expression levels of *Il-1β* and *iNos* (Fig. 3B and D). To gain more insight of the role of the CX3CL1/CX3CR1 axis on NF-κB signalling, we analysed the expression pattern of NF-κB pathway in *Cx3cr1*-deficient primary microglia. Our results show that the absence of CX3CR1 leads to a 50% of decrease in the mRNA expression of *Rela* as well as *Il-1β* and *Il-6* (Fig. 4A), without any change in *Tnf-α*, which agrees with the data depicted in Fig. 3. As the signalling pathway of NRF2 is also activated by the CX3CL1 and the promoter of *Nfe2l2* (NRF2 gene) contains a κB2 consensus sequence for NF-κB binding (Hayes & Dinkova-Kostova, 2014, Lastres-Becker et al., 2014), we assessed whether NRF2 signalling is impaired in *Cx3cr1*-deficient primary microglia. *Nfe2l2* mRNA expression levels were decreased about 50% in the absence of CX3CR1 as NRF2-dependent genes like *Nqo1, Gclc* and *Gclm* (Fig. 4B). Moreover, to determine if NRF2 activation could improve this impairment, *Cx3cr1^+/+^*and *Cx3cr1^−/−^* primary microglia were treated with sulforaphane (SFN) (15 μM, 6h), a NRF2 inducer (Jazwa, Rojo et al., 2011). Although *Cx3cr1^+/+^*microglia showed induction of *Nfe2l2, Nqo1, Gclc* and *Gclm* expression levels, *Cx3cr1^−/−^* failed to replicate this effect. These results are specific for CX3CR1-expressing microglia given that astrocytes obtained in the same purification setting did not show those effects (Suppl. Fig. 2) and exhibited SFN dependent induction.

**Figure 4:**
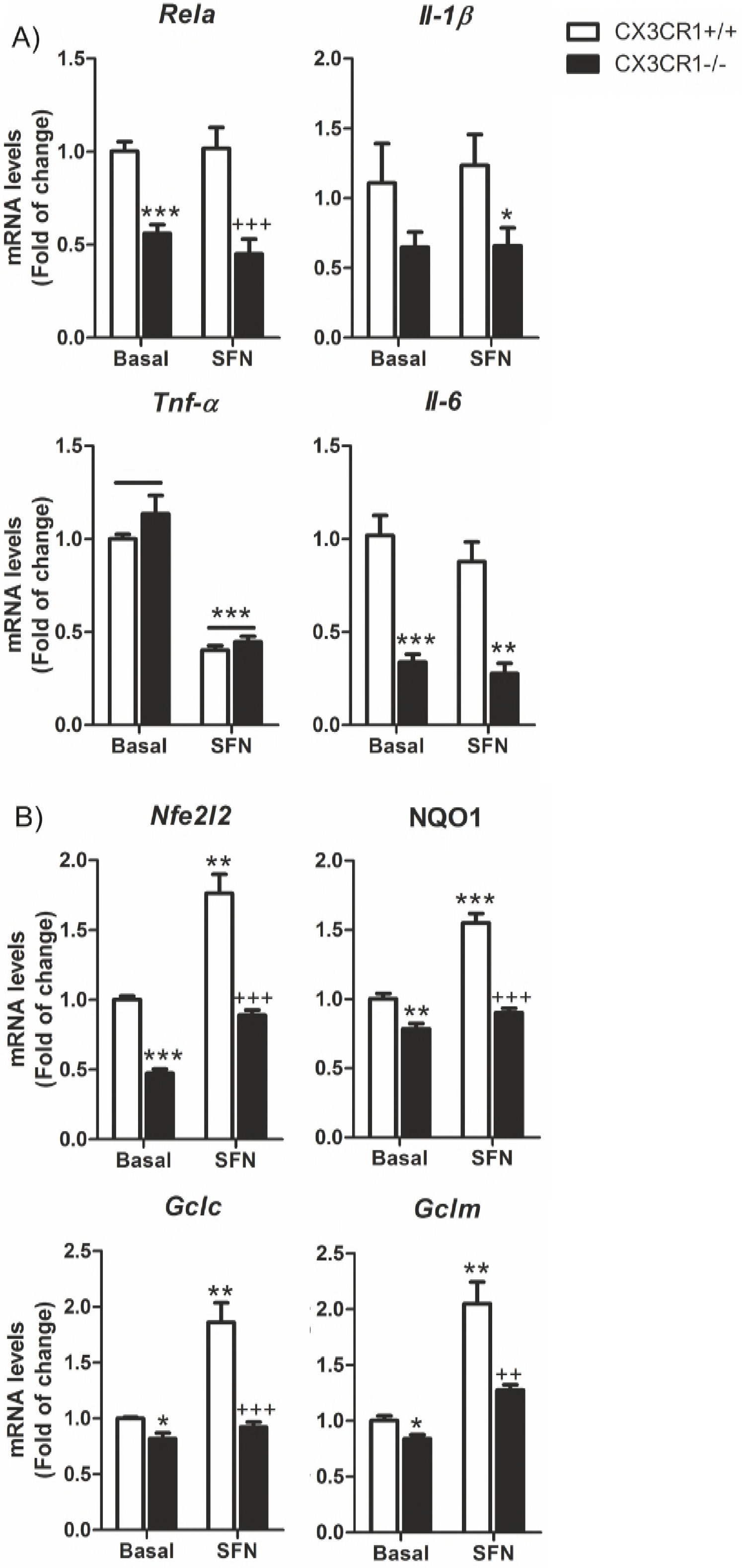
CX3CR1 receptor implications in the transcription factors NF-κB and NRF2 signalling in microglia. Primary cultures of microglia from control wild-type mice (Cx3cr1^+/+^) and Cx3cr1-knockout mice (Cx3cr1^−/−^) were incubated with SFN (15 μM, 6h). (**A**) Quantitative real-time PCR determination of messenger RNA levels of NF-κB-regulated genes coding *RelA, Il-1β, Il-6* and *Tnf-α*, respectively, normalized by *Actb* (β-Actin) messenger RNA levels. (**B**) Quantitative real-time PCR determination of messenger RNA levels of NRF2-regulated genes coding *Nfe2l2, Nqo1, Gclc* and *Gclm*, respectively, normalized by *Actb* (β-Actin) messenger RNA levels. (**C**) The promoter of *Nfe2l2* (NRF2) gene contains a κB2 sequence for NF-κB binding, modification from (Hayes & Dinkova-Kostova, 2014). Two-way ANOVA followed by Bonferroni posttest was used to assess significant differences among groups. Asterisks denote significant differences *p<0.05, **p<0.01 respect to the basal *Cx3cr1*^+/+^ group and #p < 0.05 and ##p<0.01 respect to the SFN-treated *Cx3cr1*^+/+^ group.

### CX3CR1-deficient primary microglial cells show impaired migration

Besides its role as pro- and anti-flammatory mediator, CX3CL1 has been implicated in microglial migration. In its soluble form, it acts as an extracellular chemoattractant promoting cellular migration (Paolicelli, Bisht et al., 2014), but there is no evidence about the role of CX3CR1 deficiency in microglial migration. We observed that *Cx3cr1*^−/−^ primary microglia showed impaired migration properties compared to *Cx3cr1*^+/+^microglia at basal level. This effect was more evident using a control of positive migration with serum (15% of FCS) (Fig. 5A). Moreover, the presence of CX3CL1 (100 nM, 16h) in the bottom chamber increased microglia migration of *Cx3cr1*^+/+^cells but not of cells with the *Cx3cr1^−/−^* genotype. The addition of the NRF2 inducer SFN (5 μM, 16h) did not show any significant effect in microglial migration. These data suggest that CX3CR1 has a significant role in microglial migration and can be induced by its ligand.

**Figure 5:**
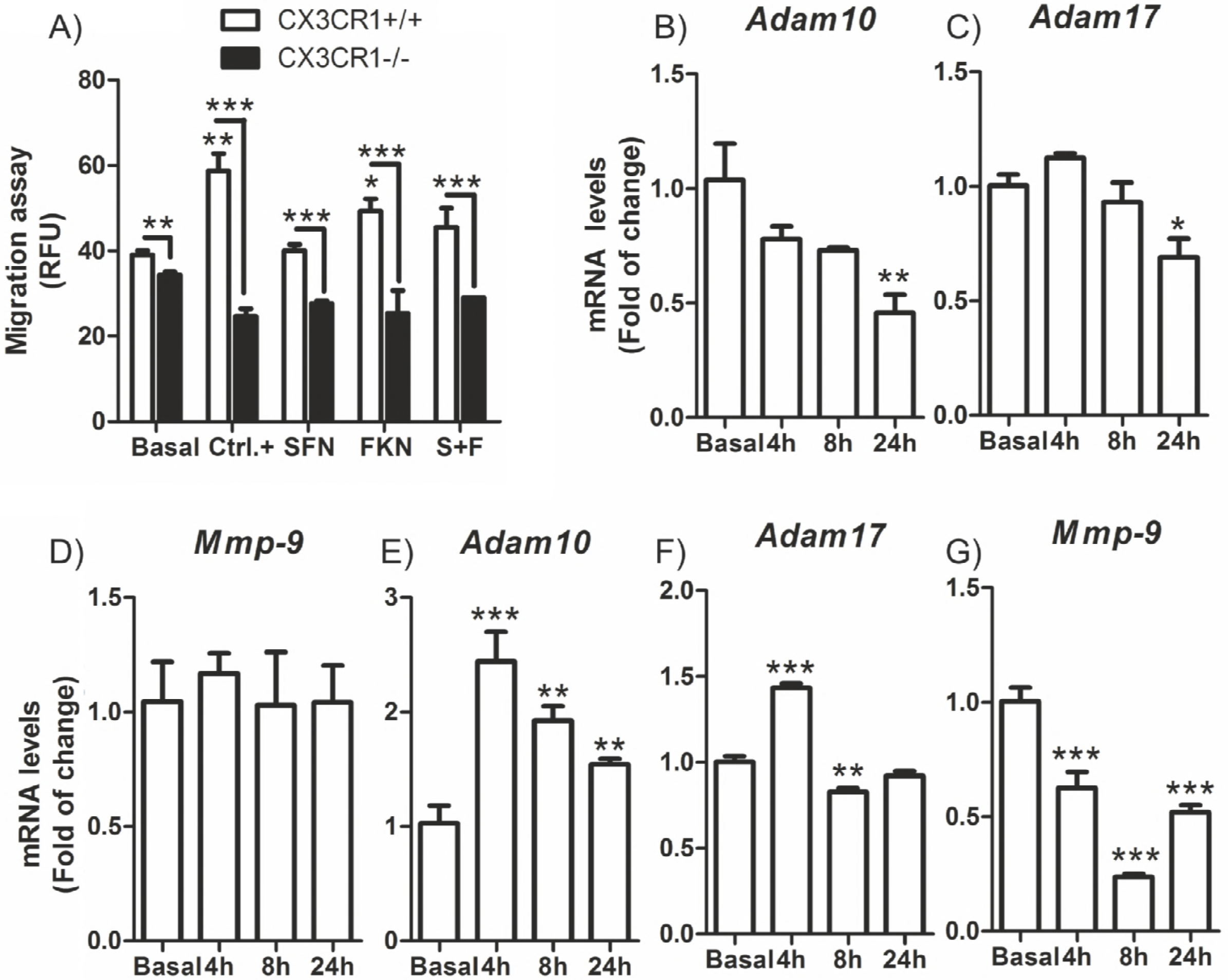
Impaired migration of Cx3cr1-deficient microglia and modulation of metalloproteinase by CX3CL1. (**A**) Motility was determined by using CytoSelect 96-well cell migration assay from primary cultures of microglia from Cx3cr1^+/+^ and Cx3cr1^−/−^ mice in accordance with the manufacturer’s instructions, described in “Materials and Methods”. Positive control: 15% FBS; SFN (5 μM); CX3CL1 (100 nM) for 16h. Values are mean±SEM (n=3, performed two times). (**B-D**) IMG cells treated with CX3CL1 (100 nM) for 4, 8 and 24h. Quantitative real-time PCR determination of messenger RNA levels of *Adam10, Adam17* and *Mmp-9*, respectively, normalized by *Actb* (β-Actin) messenger RNA levels. (**E-G**) IMG cells treated with SFN (10 μM) for 4, 8 and 24h. Quantitative real-time PCR determination of messenger RNA levels of *Adam10*, *Adam17* and *Mmp-9*, respectively, normalized by *Actb* (P-Actin) messenger RNA levels. Values are mean±SEM (n=4). Statistical analyses were performed with one-way ANOVA followed by Newman-Keuls multiple-comparison test. *p<;0.05, **p<0.01, and ***p<0.001, comparing the indicated groups.

As mentioned before, CX3CL1 can exist as a membrane-bound form (important for adhesion) and also as a cleaved product by metalloproteinase domain-containing protein 10 (ADAM10), TACE (tumor necrosis factor-α-converting enzyme)/ADAM17 and cathepsin S, depending on the cell type and the microenvironment (O’Sullivan, Gasparini et al., 2016). Furthermore, matrix metalloproteinases (MMPs) are involved in pathogenesis of neuroinflammatory diseases, while microglia are the major sources of MMPs. For example, MMP-9 contributes to inflammatory glia activation and nigrostriatal pathway degeneration in mouse models of Parkinson’s disease (Annese, Herrero et al., 2015). Hence, we investigated whether there was a possible connection between CX3CL1 and the expression of metalloproteinases along with any implication in microglia activation. Our results showed that in IMG cells, CX3CL1 treatment (100 nM) reduced mRNA levels of *Adam10* and *Adam17* after 24h, without changing *Mmp-9* levels (Fig. 5B-D). These results suggest that CX3CL1 can modulate the expression of the metalloproteinases implicated in its solubilisation. On the other hand, activation of NRF2 signalling by SFN (10 μM) induced the expression of *Adam10* and *Adam17* in a time-dependent way reaching a maximum after 4h of treatment (Fig. 5E,F). Conversely, SFN treatment decreased the expression of Mmp-9 also in a time-dependent way (Fig. 5G), suggesting an anti-inflammatory effect.

### CX3CR1-deficient microglia exhibit decreased expression of TAM receptors and phagocytosis deficiency

Phagocytosis is one of the main functions of microglial cells. TAM receptor tyrosine kinases Mer, Axl and Tyro3 regulate this microglial function (Fourgeaud, Traves et al., 2016). These receptors are essential for the phagocytosis of apoptotic cells. In the immune system, they act as pleiotropic inhibitors of the innate inflammatory response to pathogens (Lemke, 2013). Deficiencies in TAM signalling are implicated in chronic inflammatory and autoimmune disease in humans (Lastres-Becker et al., 2012, Lu & Lemke, 2001). In primary culture of microglial cells from *Cx3cr1*^+/+^and *Cx3cr1*^−/−^ mice, we observed a reduction of the expression of Axl, *Mer* and *Tyro3* mRNA levels (Fig. 6A) in *Cx3cr1*-deficient microglia. This was very similar to the reduction of the same set of genes observed in primary microglia from *Nrf2*^−/−^ mice (Lastres-Becker et al., 2012). Then, we analysed whether induction of NRF2 by SFN could restore TAM expression levels. SFN treatment (15 μM, 6h) only induced the expression of *Axl* in the same murine cells, suggesting that *Axl* promoter could possess an antioxidant response element (ARE) and be an NRF2-dependent gene. In order to test this hypothesis; we first searched the Encyclopedia of DNA Elements at UCSC (ENCODE) of the human genome (Feb. 2009) to look whether TAM receptors possess putative AREs. This database contains the experimental data from chromatin immunoprecipitation (ChIP) studies of several transcription factors, like MAFK and BACH1 which are ARE binding factors, because NFE2L2 is not included in ENCODE (Fig. 6B). We found evidence of MAFK and BACH1 consensus binding sites only in *Axl*. Then, we developed a Python-based bioinformatics analysis script (Pajares, Jiménez-Moreno et al., 2016), to compare the human consensus ARE sites from the JASPAR database26 with putative AREs in the promoter regions of AXL. We detected one ARE (relative score over 80%) in this gene. To corroborate this finding we analysed the induction of a firefly luciferase reporter containing −2376 to +7 Axl promoter (full promoter of Axl) (Mudduluru & Allgayer, 2008) by a stable mutant of NRF2, NRF2^ΔETGE^-V5, that lacks four residues (ETGE) essential for recognition by the E3 ligase complex Cul3/Keap1 or treatment with SFN (3μM for 16h). We found a dose-dependent activation of Axl reporter (Fig. 6C) by NRF2^ΔETGE^-V5 and at the same level as SFN. These data indicate that AXL is a NRF2-dependent gene. Additionally, CX3CL1 (100 nM) was able to induce Axl expression after 24h of treatment (Fig. 6D).

**Figure 6:**
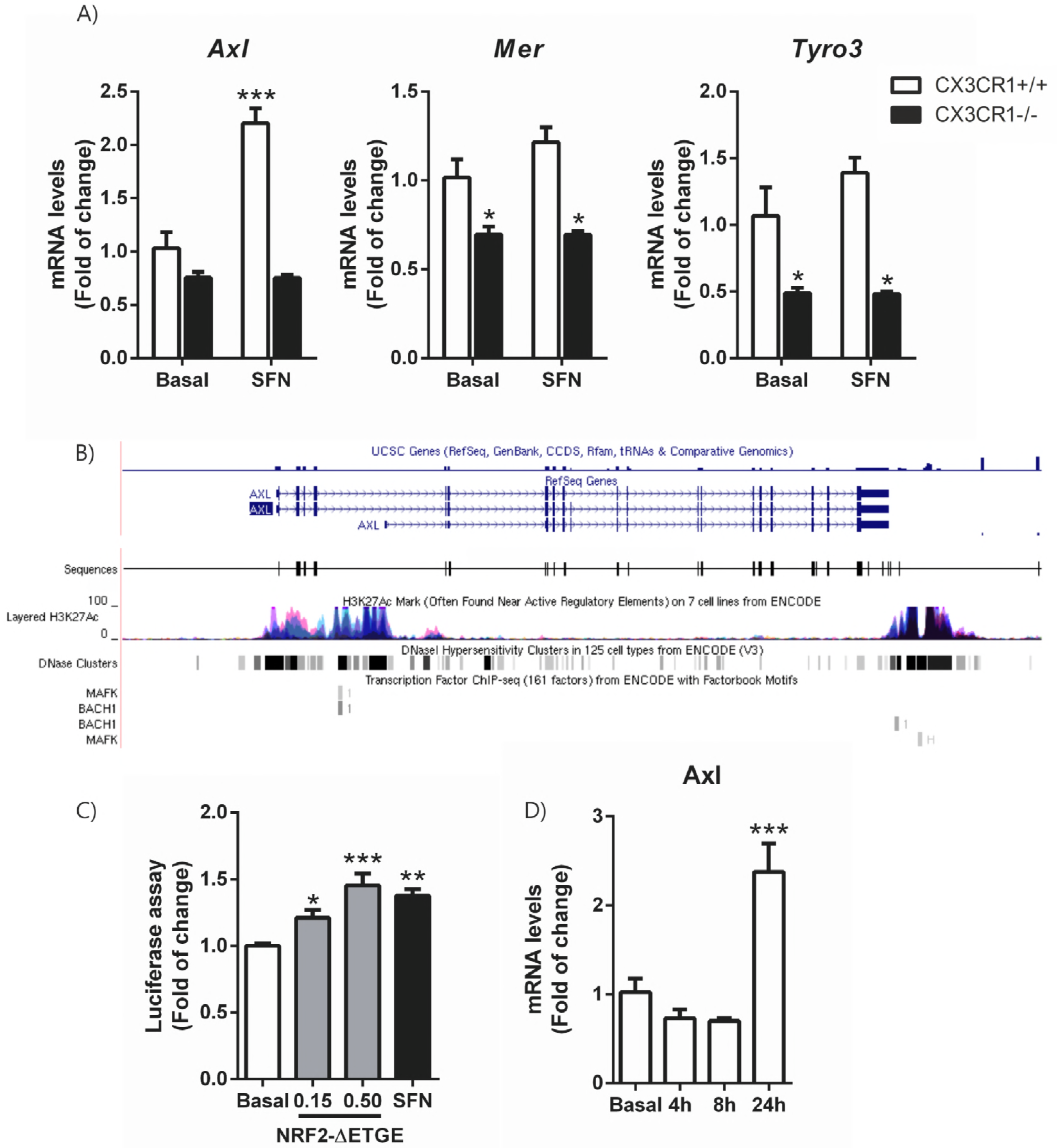
TAM receptors expression are decreased in Cx3cr1-deficient microglia. Axl could be modulated by the transcription factor NRF2. (**A**) Primary cultures of microglia from Cx3cr1^+/+^ and Cx3cr1^−/−^ mice were incubated with SFN (15 μM, 6h). Quantitative real-time PCR determination of messenger RNA levels of Axl, *Mer* and *Tyro3* respectively, normalized by *Actb* (β-Actin) messenger RNA levels. Values are mean±SEM (n=4). (**B)** To analyze the role of NFE2L2 in the transcriptional regulation of TAM receptors, we searched the Encyclopedia of DNA Elements at UCSC (ENCODE)25 of the human genome (Feb. 2009) for putative AREs. This database contains the experimental data from chromatin immunoprecipitation (ChIP) studies of several transcription factors. Although NFE2L2 is not included, we analyzed 2 other ARE binding factors, MAFK and BACH1, for which information is available. We found evidence of MAFK or BACH1 binding in the Axl gene sequence. (**C**) HEK293T cells were co-transfected with NRF2^ΔETGE^-V5 expression vector, AXL-LUC reporter, Renilla control vector and empty vector, or treated with SFN (3μM for 16h). Luciferase experiments were performed at least three times using three samples per group. (**D**) IMG cells were incubated in the presence of recombinant CX3CL1 (100 nM) for 4, 8 and 24 h and quantitative real-time PCR determination of messenger RNA for *Axl* was analysed and normalized by *Actb* (β-Actin) messenger RNA levels. Values are mean±SEM. Statistical analyses were performed with one-way ANOVA followed by Newman-Keuls multiple-comparison test: *p<0.05, *<<0.01, and ***p<0.001, comparing the indicated groups.

To get insight in the functional implications of CX3CR1 deficiency in microglia, we analysed the phagocytic capacity of *Cx3cr1*^+/+^ and *Cx3cr1*^−/−^ primary microglial murine cells at basal levels and after stimulation with CX3CL1. Lack of CX3CR1 reduces significantly the phagocytic capacity of microglial cells (Fig. 7). Besides, CX3CL1 is able to increase phagocytosis in *Cx3cr1*^+/+^ microglia but not in *Cx3cr1*^−/−^ cells, indicating the relevance of CX3CR1 in one of the main function of microglia. Interestingly, these results are very similar to those obtained in the microglia deficient for NRF2 (Lastres-Becker et al., 2012).

**Figure 7:**
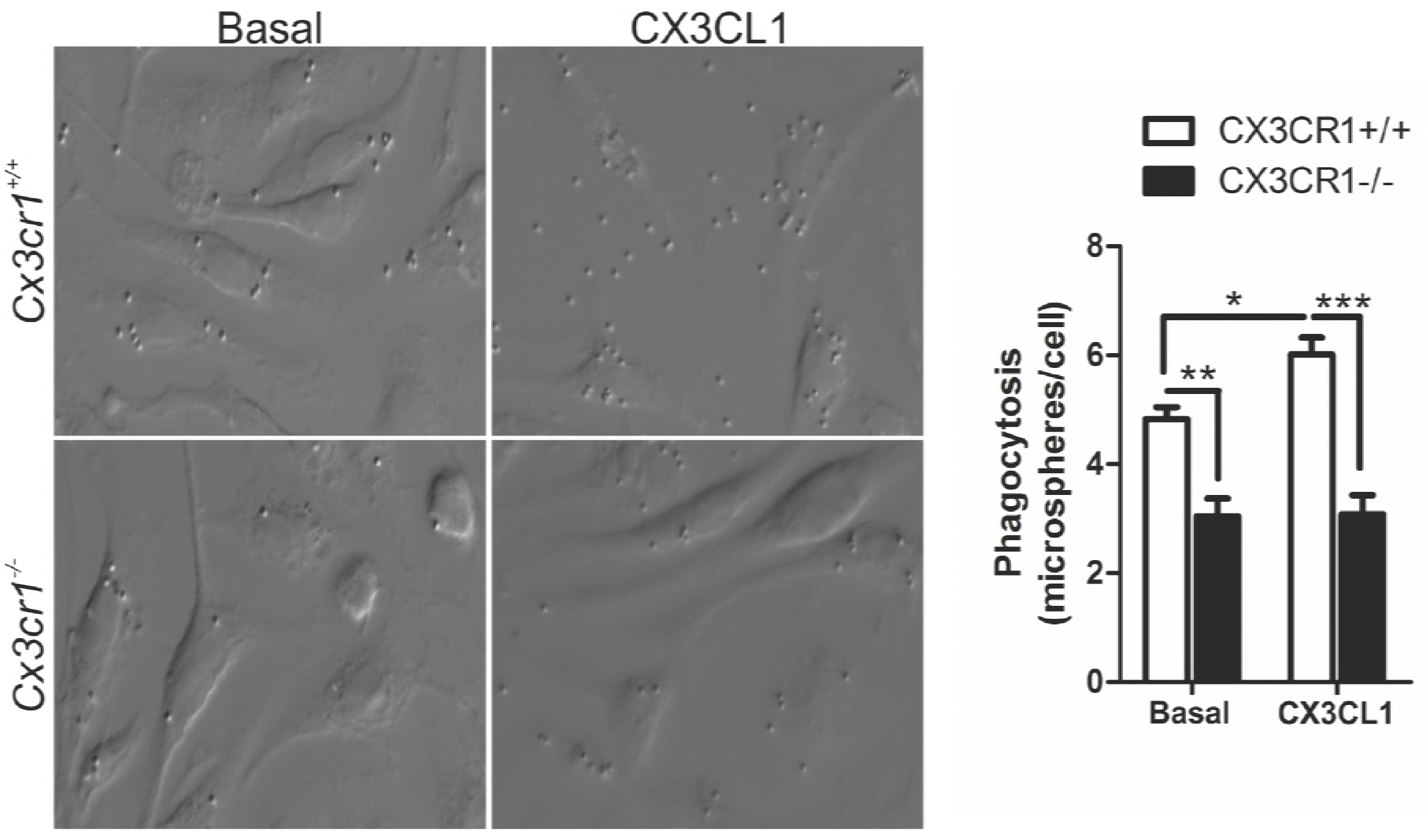
Effect of CX3CR1-deficiency on the phagocytic response. Microglia from Cx3cr1^+/+^ or Cx3cr1^−/−^ mice were incubated with fluorescent microspheres in the absence or presence of 100 nM of CX3CL1 for 2 h. Phagocytic efficiency was calculated as a number of microspheres per cell. One-way ANOVA followed by Newman-Keuls test was used to assess differences among groups. Asterisks denote significant differences: *p<0.05, **p<0.01, ***p<0.001 compared with the indicated groups.

### SFN treatment reverses astrogliosis but not microgliosis in the *Cx3cr1*^−/−^ mice in the rAAV-TAU^P301L^ mouse model

Previous work from our group (Lastres-Becker et al., 2014) showed the relevance of the CX3CL1/CX3CR1/NRF2 axis in a tauopathy mouse model. In this model, which is based on stereotaxic delivery in hippocampus of an adeno-associated viral vector for expression of TAU^P301L^, we confirmed that TAU-injured neurons express CX3CL1, and that NRF2-and CX3CR1-knockout mice not only do not express heme oxygenase 1 (HO1) in microglia but also exhibited exacerbated microgliosis and astrogliosis. Considering that in CX3CR1-deficient mice there is a reduction of the NRF2 axis in microglial cells, we evaluated the effect of an inducer of NRF2 as a possible modulator of this effect with putative implications the neuroinflammatory associated to tauopathies. Moreover, it has been reported that NRF2 induction has neuroprotective effects in several neurodegenerative mouse models (Jazwa et al., 2011, Lastres-Becker et al., 2016, Petrillo, Piermarini et al., 2017, Phillips & Fox, 2013) as well as in tauopathies (Cuadrado et al., 2018). Thus, the effects of SFN on neuroinflammation were examined in *Cx3cr1*^+/+^and *Cx3cr1*^−/−^ mice stereotaxically injected in the right hippocampus with AAV-TAU^P301L^ and treated daily with SFN (50mg/kg, i.p) during three weeks. Overexpression of TAU^P301L^ induced microgliosis at the ipsilateral side of injection (Suppl. Fig. 3) in *Cx3cri*^+/+^ mice and was slightly exacerbated in *Cx3cr1*^−/−^ mice (Suppl. Fig 3 and Fig. 8A, C and E). SFN treatment could attenuate significantly microgliosis in *Cx3cri*^+/+^but not in *Cx3cr1*^−/−^ mice (Fig. 8B, D and E). We analysed HO1 expression to determine whether SFN was able to induce NRF2 signalling in microglial cells, because the commercially available NRF2 antibodies are in general not specific (Lau, Tian et al., 2013). SFN treatment induced HO1 expression in microglial cells (colocalization of IBA1 and HO1) and astrocytes of the contralateral hippocampus (Fig. 8A and B) from *Cx3cr1*^+/+^ mice (Fig.9). Furthermore, TAUP301L overexpression increased the number of microglial cells positive for HO1 (Fig. 8A). SFN treatment did not show any HO1-expression alterations (Fig. 8B). On the other hand, in *Cx3cri*^−/−^ mice, microglial cells did not express HO1 either at basal levels (Fig. 8C) or after SFN treatment (Fig. 8D). Additionally, TAUP301L overexpression did not induce the expression of HO1 in microglial cells (neither VEH nor SFN treatment did). These results indicate that SFN treatment is not able to induce NRF2/HO1 signalling in *Cx3cr1*^−/−^ microglia and therefore it cannot modulate the microgliosis induced by TAUP301L overexpression. Whereas SFN treatment reversed astrogliosis in both genotypes (Fig. 9), indicating that NRF2 activation was sufficient to modulate reactive astrocytosis. This was corroborated by the observation that astrocytes were able to express HO1 (Fig. 9).

**Figure 8:**
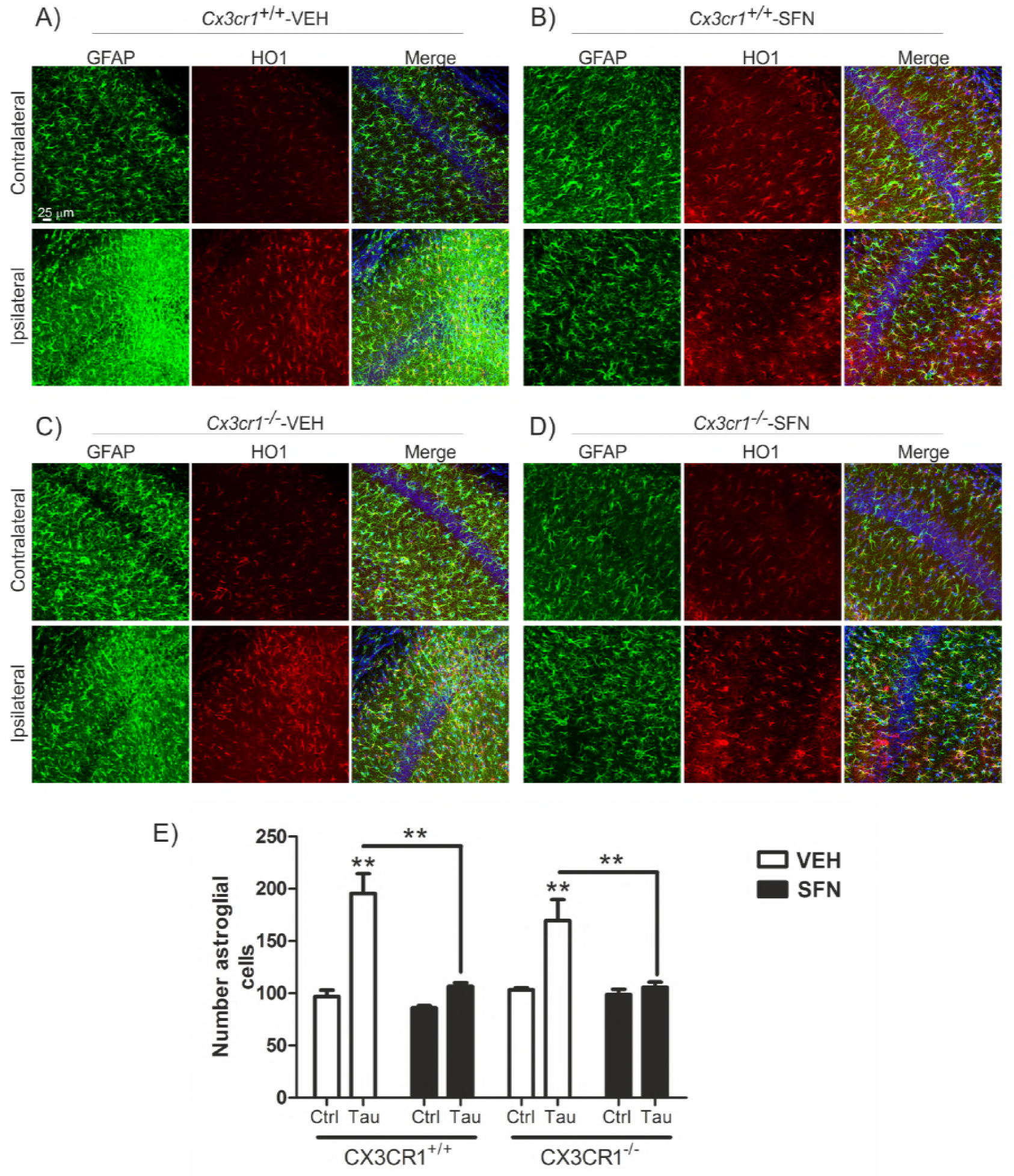
SFN treatment attenuates TAU^P301L^-induced astrogliosis. Photographs show the astrocyte marker GFAP (green) and HO1 (red) as a NRF2 reporter gene, in 30 μm-thick sections of hippocampus from mice with the genotypes (**A**) *Cx3cr1*^+/+^-VEH, (**B**) Cx3cr1^+/+^-SFN, (**C**) *Cx3cr1*^−/−^-VEH and (**D**) Cx3cr1^+/+^-SFN. (**E**) Stereological quantification of the number of astrocytes in the control side and the TAU^P301L^ expressing side of all the experimental groups. Differences among groups were assessed by two-way ANOVA followed by Bonferroni’s test. Asterisks denote significant differences **p<0.01, comparing the indicated groups.

**Figure 9:**
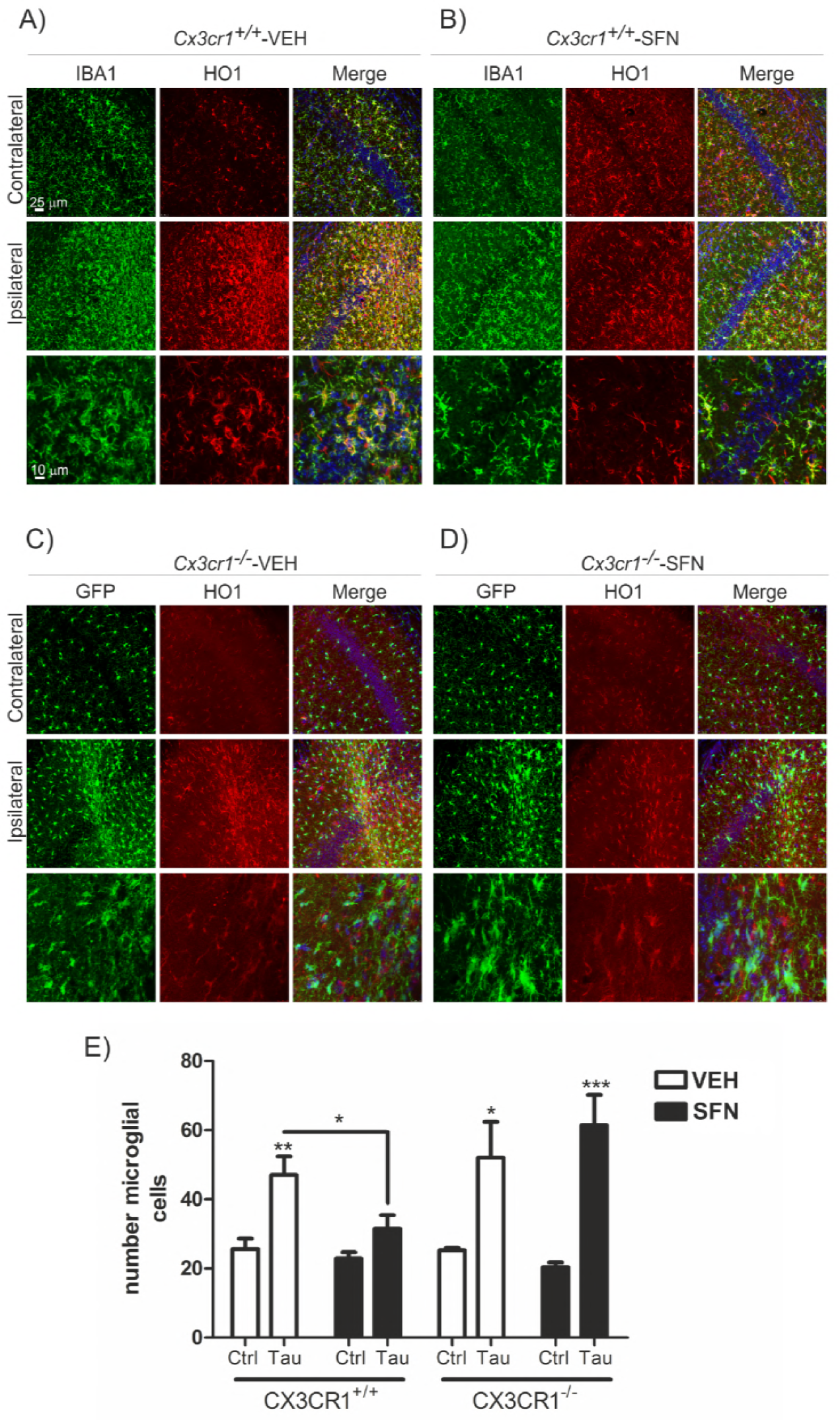
CX3CR1 receptor is required to induce HO1 in microglia in response to TAU^*P301L*^ expression and SFN treatment. Photographs show the microglial marker IBA1 (green) and HO1 (red) as a NRF2 reporter gene, in 30 μm-thick sections of hippocampus from mice with the genotypes (**A**) *Cx3cr1*^+/+^-VEH, (**B**) Cx3cr1^+/+^-SFN, (**C**) *Cx3cr1*^−/−^-VEH and (**D**) *Cx3cr1*^−/−^-SFN. Microglia from *Cx3cr1*^−/−^ mice, stained with anti-GFP antibody (green), do not express HO1 (red) in response to TAU^P301L^. (**E**) Stereological quantification of the number of microglial cells in the control side and the TAU^P301L^ expressing side of all experimental groups. Two-way ANOVA followed by Bonferroni post-test was used to assess significant differences among groups. Asterisks denote significant differences *p<0.05, **p<0.01 and ***p<0.001, comparing the indicated groups.

## DISCUSSION

Neurodegenerative diseases such as Alzheimer’s disease and other tauopathies are among the most burdensome health concerns in societies with pronounced aging tendencies. Although neurons are essential targets in neurodegeneration, there are other environmental factors that contribute to neurodegeneration such as neuroinflammation, which involves other cell types of the brain. In this context, the study of non-cell-autonomous neurodegenerative process would open new windows to understand how neurodegeneration could be promoted by local neuroinflammation. Microglia play an essential role in development, homeostasis, synapse modulation, phagocytosis and neurogenesis, indicating that impairments in the crosstalk between neurons and microglia are determinant in neurodegenerative processes. Fractalkine (CX3CL1) and its cognate receptor (CX3CR1) are key factors in this crosstalk, although there is great controversy on whether it promotes proinflammation or has any anti-inflammatory effect (Bhaskar, Konerth et al., 2010, Jones, Beamer et al., 2010, Lauro et al., 2015). In this study, we showed that treatment of immortalized microglial cells (IMG) with CX3CL1 induced NF-κB-p65 and pro-inflammatory cytokines expression (Fig. 1). Phospho-kinase assay proteome profiler (in IMG and primary microglial cells) indicates that CX3CL1 triggers several kinase pathways such as ERK1/2, p38 and JNK that converge to MSK1-dependent NF-κB signalling activation (Fig. 2-3 and Suppl. Figure 4). These results were corroborated in siMSK1 microglial cells, which displayed impaired NF-κB signalling after CX3CL1 treatment. On the other hand, CX3CR1-deficient primary microglial cells present impaired levels of NF-κB mRNA expression and decreased anti-inflammatory NRF2 signalling, suggesting a dual role of the CX3CL1/CX3CR1 axis in neuroinflammation (Fig. 4). In addition, CX3CR1-deficient primary microglial cells show impaired migration at basal levels and under CX3CL1 stimulation (Fig. 5). Microglia lacking CX3CR1 expression exhibit decreased mRNA levels of TAM receptors (TYRO3, AXL and MER), which functionally results in deficiency of phagocytosis (Fig. 6-7) (to summarize all the molecular pathways see Suppl. Fig. 4). These observations are supported by the results from Dr. Jesus Avila’s laboratory, which demonstrated that the CX3CL1/CX3CR1 axis played a key role in the phagocytosis of TAU by microglia *in vitro* and *in vivo* and that this was also affected as AD progressed. Moreover, they found a novel mechanism of TAU internalization by microglia through direct binding to CX3CR1 (Bolos, Llorens-Martin et al., 2017). Taken together, these data suggest that CX3CR1 could be a critical key factor in microglial function and tauopathies progression.

Interestingly, in relation to the CX3CR1 receptor, 2 polymorphisms named V249I and T280M have been described, which induce a low availability of receptors on the cell surface or have a reduced receptor affinity for CX3CL1, respectively. It has been shown that CX3CR1 is a gene involved in the modification of survival and progression in amyotrophic lateral sclerosis (Lopez-Lopez et al. 2014), and that there is an association of the variant CX3CR1-V249I with a progression of neurofibrillary pathology in late-onset Alzheimer’s disease (Lopez-Lopez et al. 2017). These studies indicate that polymorphisms in CX3CR1 could modulate neurodegenerative progression (Suppl. Figure 5), as well polymorphisms in NRF2 (Cho, Marzec et al., 2015). Therefore, our study is pioneer in the description and analysis of the molecular pathways involved in the activation of the CX3CL1/CX3CR1 axis and their implications in the treatment of taupathies.

Regarding the TAM receptors, which regulate cell migration, survival, phagocytosis and clearance of metabolic products or cell debris (Shafit-Zagardo, Gruber et al., 2018), we found that CX3CR1 deficiency decreased TAM mRNA expression levels, and that only Ax1 could be modulated by the activation of NRF2 by SFN. Interestingly, these results are comparable to those obtained in primary microglia of NRF2-deficient mice (Lastres-Becker et al., 2014, Lastres-Becker et al., 2012), indicating a crosstalk between CX3CR1/NRF2/TAM receptors and also its implication in phagocytosis. Therefore, to determine whether TAM receptors genes could be NRF2-dependent we screened the chromatin immunoprecipitation database ENCODE for 2 proteins, MAFK and BACH1 that bind the NFE2L2-regulated enhancer antioxidant response element (ARE). Using a script generated from the JASPAR’s consensus ARE sequence, we identified only 1 putative ARE in the Axl gene (Fig. 6). This result was corroborated using a luciferase reporter Axl-LUC, which contains the full promoter of Axl. Different concentrations of a stable mutant of NRF2, NRF2ΔETGE-V5, that lacks four residues (ETGE) essential for recognition by the E3 ligase complex Cul3/Keap1, induce Axl in a dose-dependent way and to a similar extent as SFN. It has been shown that Gas6-Axl signalling plays an important role in maintaining axonal integrity in addition to regulating and reducing CNS inflammation that cannot be compensated for by ProS1/Tyro3/MerTK signalling (Ray, DuBois et al., 2017). Hence, modulation of Axl by NRF2-inducers could become a new potential target for therapeutic intervention in CNS diseases.

Finally, the effects of SFN were examined on neuroinflammation in *Cx3cr1*^+/+^ and *Cx3cr1*^−/−^ mice stereotaxically injected in the right hippocampus with AAV-TAU^P301L^ and treated daily with SFN (50mg/kg, i.p) during three weeks. Whereas SFN treatment reversed astrogliosis in both genotypes (Fig. 8), at the microglia level we did not see any improvement in the *Cx3cr1*^−/−^ mice (Fig. 9), obtaining in this way similar effects as in NRF2-deficient mice. These findings suggest that the CX3CR1-NRF2 axis activation is essential for the modulation of microglial activation associated with tauopathies, and that the associated polymorphisms of CX3CR1 (Suppl. Fig. 4) must be taken into account in the design of pharmacological strategies aimed to the treatment of these diseases.

## MATERIAL AND METHODS

### Cell culture

Primary astrocytes and microglia were prepared from neonatal (P0-P2) mouse cortex from *Cx3cr12*^+/+^ and *Cx3cr1*^−/−^ and grown and isolated as described in (Lastres-Becker et al., 2014). Briefly, neonatal (P0-P2) mouse cortex were mechanically dissociated and the cells were seeded onto 75 cm2 flasks in DMEM:F12 supplemented with 10% FCS and penicillin/streptomycin. After 2 weeks in culture, flasks were trypsinized and separated using CD11b MicroBeads for magnetic cell sorting (MACS Miltenyi Biotec, Germany). Microglial and astroglial cultures were at least 99% pure, as judged by immunocytochemical criteria. Medium was changed to Dulbecco’s Modified Eagle Medium:F12 (DMEM:F12) serum-free without antibiotics 16 h before treatment. Immortalized microglial cell line (IMG) isolated from the brains of adult mice, were purchased from Kerafast Inc., and were grown in Dulbecco’s Modified Eagle Medium (DMEM) supplemented with 10% fetal bovine serum, 1% penicillin/streptomycin and 2 mM L-glutamine, in 5% CO_2_ at 37°C, 50 % relative humidity. Medium was changed to serum-free DMEM without antibiotics 16 h before treatments.

### Preparation of nuclear and cytosolic extracts

IMG cells were seeded in p100 plates (1.5 × 10^6^ cells/plate). IMG cells were treated with FKN (100 nM) and samples collected at different time points. Cytosolic and nuclear fractions were prepared as described previously (Rojo, Salinas et al., 2004). Briefly, cells were washed with cold PBS and harvested by centrifugation at 1100 rpm for 10 min. The cell pellet was resuspended in 3 pellet volumes of cold buffer A (20 mM HEPES, pH 7.0, 0.15 mM EDTA, 0.015 mM EGTA, 10 mM KCl, 1% Nonidet P-40, 1 mM phenylmethylsulfonyl fluoride, 20 mM NaF, 1 mM sodium pyrophosphate, 1 mM sodium orthovanadate, 1 μg/ml leupeptin) and incubated in ice for 30 min. Then the homogenate was centrifuged at 500 *g* for 5 min. The supernatants were taken as the cytosolic fraction. The nuclear pellet was resuspended in 5 volumes of cold buffer B (10 mM HEPES, pH 8.0, 0.1 mM EDTA, 0.1 mM NaCl, 25% glycerol, 1 mM phenylmethylsulfonyl fluoride, 20 mM NaF, 1 mM sodium pyrophosphate, 1 mM sodium orthovanadate, 1 μg/ml leupeptin). After centrifugation in the same conditions indicated above, the nuclei were resuspended in loading buffer containing 0.5% SDS. The cytosolic and nuclear fractions were resolved in SDS-PAGE and immunoblotted with the indicated antibodies.

### Immunoblotting

Whole cell lysates were prepared in RIPA-Buffer (25 mM Tris-HCl pH 7.6, 150 mM NaCl, 1 mM EGTA, 1% Igepal, 1% sodium deoxycholate, 0.1 % SDS, 1 mM PSMF, 1 mM Na3VO4, 1 mM NaF, 1 μg/ml aprotinin, 1 μg/ml leupeptin and 1 μg/ml pepstatin). Whole cell lysates, cytosolic and nuclear fractions containing 25 μg of whole proteins from IMG-treated cells were loaded for SDS-PAGE electrophoresis. Immunoblots were performed as described in Cuadrado et al., 2014. The primary antibodies used are described in Supplementary Table S1.

*Analysis of mRNA levels by quantitative real-time PCR*. Total RNA extraction, reverse transcription, and quantitative polymerase chain reaction (PCR) were done as detailed in previous articles (Lastres-Becker et al., 2014). Primer sequences are shown in Supplementary Table S2. Data analysis was based on the ΔΔCT method with normalization of the raw data to housekeeping genes (Applied Biosystems). All PCRs were performed in triplicates.

### Phospho-kinase assay proteome profiler

Primary microglia from *Cx3cr1*^+/+^ and *Cx3cr1*^−/−^ mice were collected as described above and 150.000 cells were plated on p60 to 80% of confluency for 16h. IMG were seeded at 300.000 cells/p60. Then, the medium was replaced with serum-free DMEM without antibiotics for 24 h before FKN (100 nM) treatment for 1h. Cells were then lysed and total proteins were extracted with lysis buffer supplemented by the kit. According to the manufacturer’s protocol, 200 μg cell lysate was incubated with each human phospho-kinase array (R&D Systems, Minneapolis, MN). Cell lysates were diluted and incubated overnight with nitrocellulose membranes in which capture and control antibodies against 43 different kinases and transcription factors, have been spotted in duplicate. The arrays were washed to remove unbound proteins and were incubated with a cocktail of biotinylated detection antibodies. Streptavidin-HRP and chemiluminescent detection reagents were applied and a signal was produced at each capture spot corresponding to the amount of phosphorylated protein bound. Pixel densities on developed X-ray film were collected and analyzed by ImageJ.

### siRNA assays

siRNA used to knock down mouse MSK1 (ON-TARGETplus Mouse Rps6ka5 (73086) siRNA-SMARTpool, Dharmacon, Catalog #L-040751-00-0005), and control scrambled siRNA sequence (Silencer® Select Negative Control siRNA #1, Ambion Cat#: 4390843). IMG cells were seeded at 500.000 cells/p60. Then, cells were transfected with 25 nM siRNAs following DharmaFECT General Transfection Protocol (Dharmacon). After 24 h, cells were transfected with 12.5 nM siRNAs and after 24h used for experiments and harvested for analysis.

### Migration assay

Cell migration was assayed using the CytoSelect 96-well Cell Migration Assay according to the manufacturer’s instruction (Cell Biolabs, Cambridge, UK). In brief, primary microglia from *Cx3cr1*^+/+^ and *Cx3cr1*^−/−^ mice were collected as described above and were resuspended in serum-free DMEM containing 0.1% bovine serum albumin (BSA). As positive control medium containing 15% fetal bovine serum was added to the bottom chamber (100 μl per well). Primary microglial cells were added to the top insert and cells were incubated at 37°C for 16 h before lysis of migratory cells and quantified using CyQuant^®^ GR Fluorescent Dye, using a fluorescence plate reader with excitation at 480 nm and emission at 520 nm.

### Luciferase asays

Transient transfections of HEK293T cells were performed with the expression vectors pGL3_AxlP1-LUC (gift of Prof. Dr H. Allgayer, Department of Experimental Surgery-Cancer Metastasis, Medical Faculty, Ruprecht-Karls-University Heidelberg, Germany). pTK-Renilla was used as an internal control vector (Promega). Luciferase assays were performed as described in Cuadrado et al., 2014.

### Bioinformatics analysis

A putative antioxidant response element (ARE) in Axl gene promoters was identified in The Encyclopedia of DNA Elements at UCSC (ENCODE)25 for the human genome (Feb. 2009) taking as reference the available information from chromatin immunoprecipitation (ChIP) of ARE binding factors MAFK and BACH1. The putative MAFK and BACH1 binding regions were localized in 200- to 400-base pair-long DNase-sensitive and H3K27Ac-rich regions, i.e., most likely regulatory promoter regions. In addition, a frequency matrix of the consensus ARE sequence based on the JASPAR database26 was converted to a position-specific scoring matrix (PSSM) by turning the frequencies into scores through the log(2) [odd-ratio (odd ratio: observed frequency/expected frequency)]. One unit was added to each frequency to avoid log(0). Then a script was generated with the Python 3.4 program to scan the promoter sequences with candidate AREs retrieved from ENCODE with the PSSM. The max score was calculated by adding the independent scores for each of the 11 base pairs of the consensus ARE sequence with the PSSM. The relative score (scorerelative) was calculated from this max score (score of the sequencemax) as: scorerelative = (score of the sequencemax-scoremin possible)/(scoremax possible - coremin possible). The min possible score (scoremin possible) is calculated as the lowest possible number obtained for a sequence from the PSSM and the max possible score (scoremax possible) is the highest possible score that can be obtained. We considered putative ARE sequences those with a scorerelative over 80%, which is a commonly used threshold for the computational framework for transcription factor binding site/TFBS analyses using PSSM.

### Phagocytosis assay

Primary microglia from *Cx3cr1*^+/+^ and *Cx3cr1*^−/−^ mice were collected as described above and 150.000 cells were plated on coverslips for 16 h. Then, the medium was replaced with serum-free DMEM:F12 without antibiotics for 24 h before adding fluorescent microspheres (150 microspheres per cell) (FluoSpheres polystyrene microspheres, Invitrogen) and FKN (100 nM) and incubating for 2 h. Then, cells were washed with PBS, fixed with 4% paraformaldehyde, and stained with DAPI. The images were captured using 90i Nikon microscope (Nikon, Montreal, QC, Canada) at 40X.

### Animals and treatments

Colonies of *Cx3cr1*^−/−^ (B6.129P-Cx3cr1^tm1Litt^/J) mice and *Cx3cr1*^+/+^ littermates were obtained from Jackson Laboratory, Bar Harbor, ME (Jung, Aliberti et al., 2000). Each experimental group comprised 5-8 animals. Recombinant AAV vectors of hybrid serotype 1/2 express mutant hTAU^P301L^ under control of the human synapsin 1 gene promoter and were used as described (Jaworski, Dewachter et al., 2009). Surgical procedures and unilateral intracerebral injection of viral particles into the right hemisphere were performed as described before (Lastres-Becker et al., 2014). In brief, 2 μL of viral suspension containing 10E8 t.u. were injected at the stereotaxic coordinates-1.94 mm posterior, −1.4 mm lateral, and −1.8 mm ventral relative to bregma. All experiments were performed by certified researchers according to regional, national, and European regulations concerning animal welfare and animal experimentation, and were authorized by the Ethics Committee for Research of the Universidad Autónoma de Madrid and the Comunidad Autónoma de Madrid, Spain, with Ref PROEX 279/14, following institutional, Spanish and European guidelines (Boletín Oficial del Estado (BOE) of 18 March 1988 and 86/609/EEC, 2003/65/EC European Council Directives). SFN (50 mg/kg) (LKT Laboratories, St. Paul, MN) was prepared in saline solution just before use and i.p. injected. We did not detect significant weight loss, hair loss or other gross alterations in the SFN-treated mice either in the 3-weeks administration every day.

### Immunofluorescence on mouse tissues

The protocol was previously described (Lastres-Becker et al., 2012). Primary antibodies are described in Supplementary Table S1. Secondary antibodies were: Alexa Fluor 546 goat anti-mouse, Alexa 546 goat antirabbit and Alexa Fluor 488 goat anti-mouse (1:500, Life technologies, Madrid, Spain). Control sections were treated following identical protocols but omitting the primary antibody.

### Statistical analyses

Data are presented as mean ± SEM. To determine the statistical test to be used, we employed GraphPad Instat 3, which includes the analysis of the data to normal distribution via the Kolmogorov-Smirnov test. In addition, statistical assessments of differences between groups were analysed (GraphPad Prism 5, San Diego, CA) by unpaired Student’s t-tests when normal distribution and equal variances were fulfilled, or by the non-parametric Mann-Whitney test. One and twoway ANOVA with *post hoc* Newman-Keuls test or Bonferroni’s test were used, as appropriate.

## Acknowledgements

This work was supported by grants from the Spanish Ministry of Economy and Competitiveness (Grants refs. SAF2016-76520-R).

## Author contributions

ILB contributed to conception and design of the study. SK, generation of AAV6-TAU(P301L) viruses. SCS, AJGY and ILB acquisition and analysis of data. ILB contributed to drafting a significant portion of the manuscript and figures.

## FIGURE LEGENDS

**Supplementary Figure 1:**
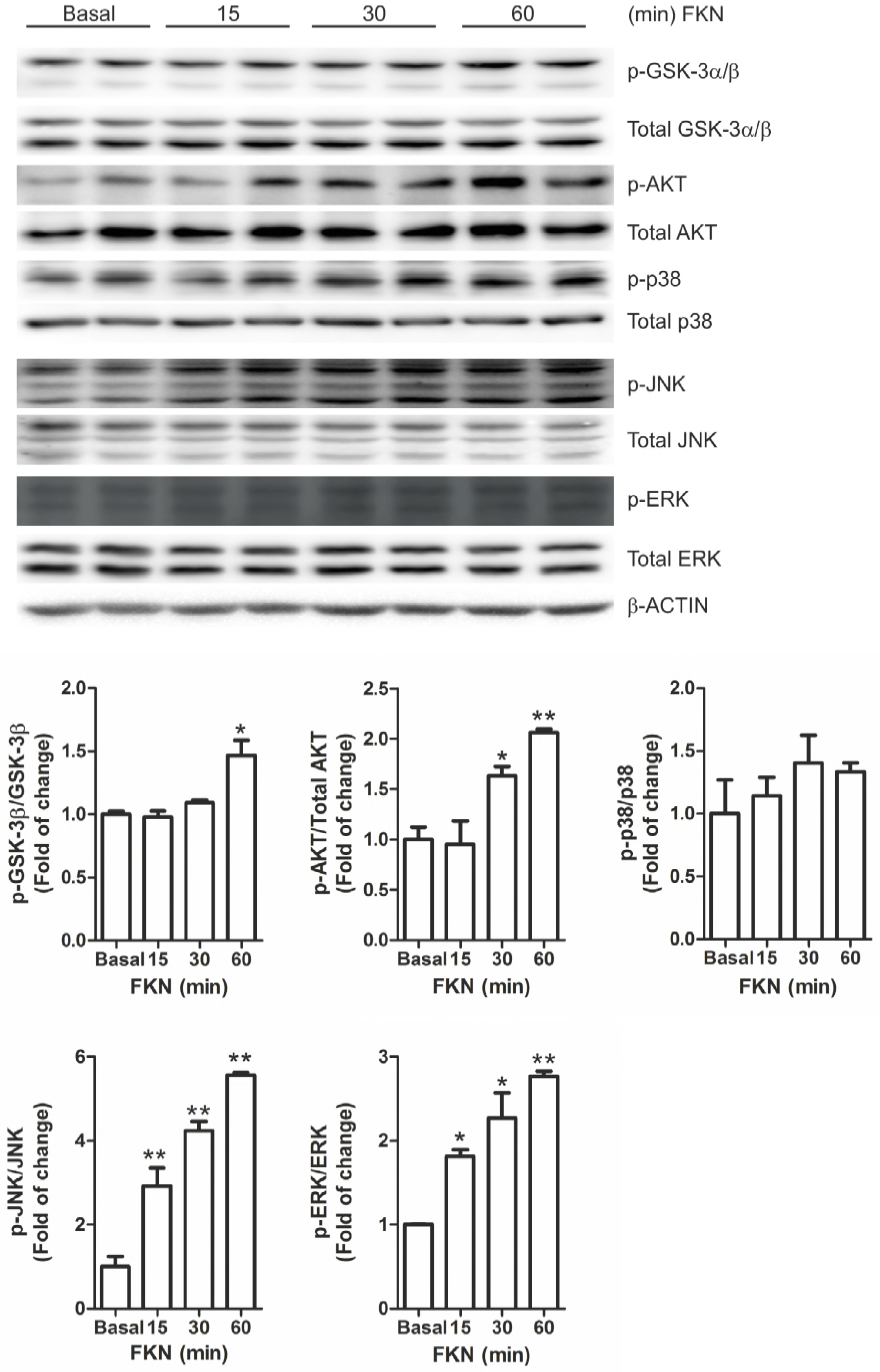
CX3CL1 activates MAPK and PI3K/AKT/GSK-3 signalling pathways. IMG cells were incubated in the presence of recombinant CX3CL1 (100 nM) for 15, 30 and 60 min, as indicated. (**A**) Analysis of total cell lysates in immunoblots: p-GSK-3α/β vs total GSK-3α/β levels; β-AKT vs total AKT levels; p-p38 vs total p38 levels; p-JNK vs total JNK levels and p-ERK vs total ERK levels. Densitometric quantification of representative blots is shown. Asterisks denote significant differences *p<0.05 and **p<0.01, comparing the indicated groups with the basal condition according to a one-way ANOVA followed by Newman-Keuls post-test.

**Supplementary Figure 2:**
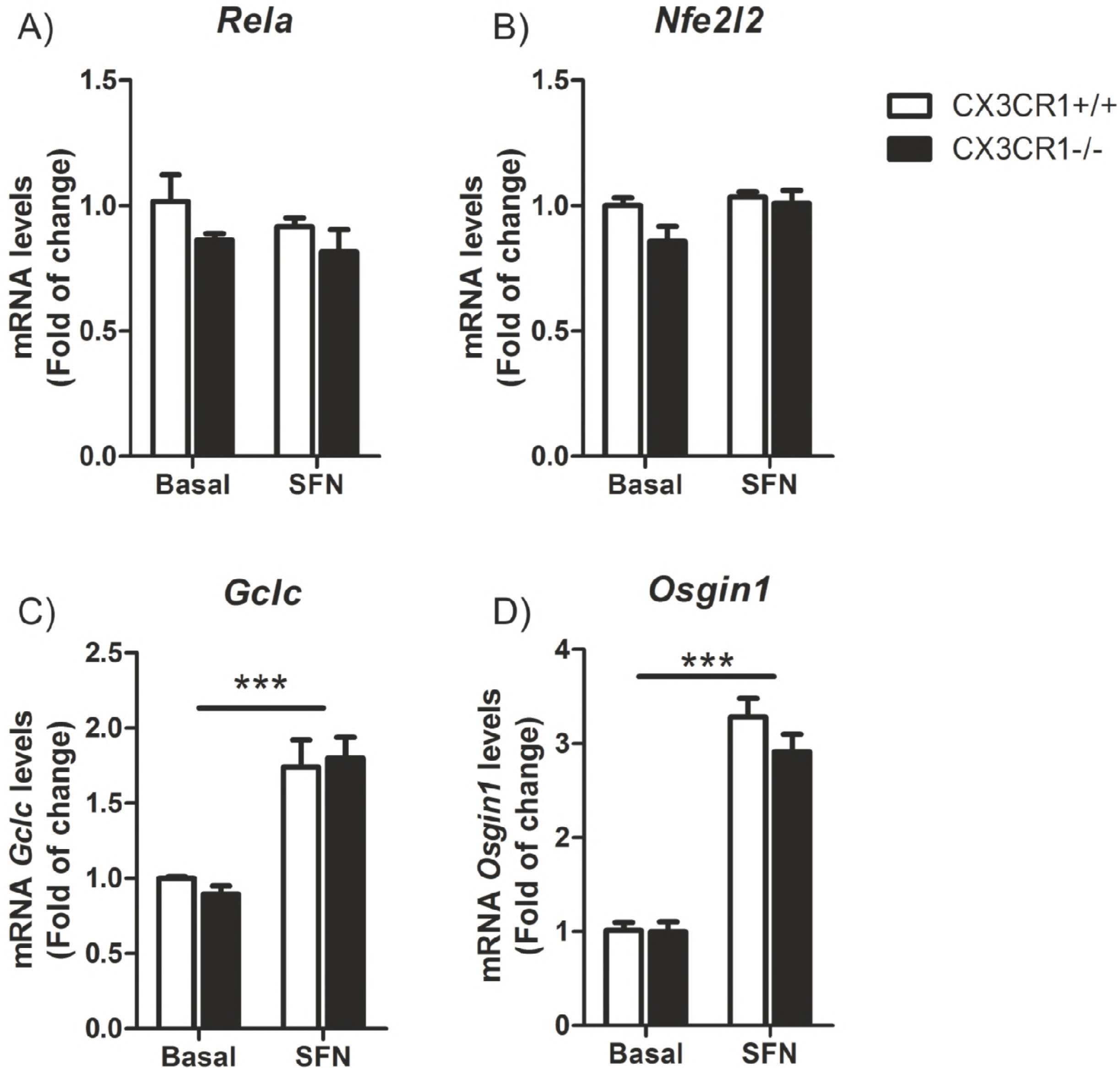
Astrocytes from Cx3cr1^−/−^ mice do not shown any alterations in the transcription factors NF-κB and NRF2 signalling. Primary cultures of astrocytes from *Cx3cr1^+/+^* and *Cx3cr1^−/−^* mice were incubated with SFN (15 μM, 6h). Quantitative real-time PCR determination of messenger RNA levels of NF-κB-regulated genes coding *RelA*, and NRF2-regulated genes coding *Nfe2l2, Nqo1, Gclc* and *Osgin1*, respectively, normalized by *Actb* (β-Actin) messenger RNA levels. Two-way ANOVA followed by Bonferroni post-test was used to assess significant differences among groups. Asterisks denote significant differences ***p<0.001 respect to the basal *Cx3cr1*^+/+^ group.

**Supplementary Figure 3:**
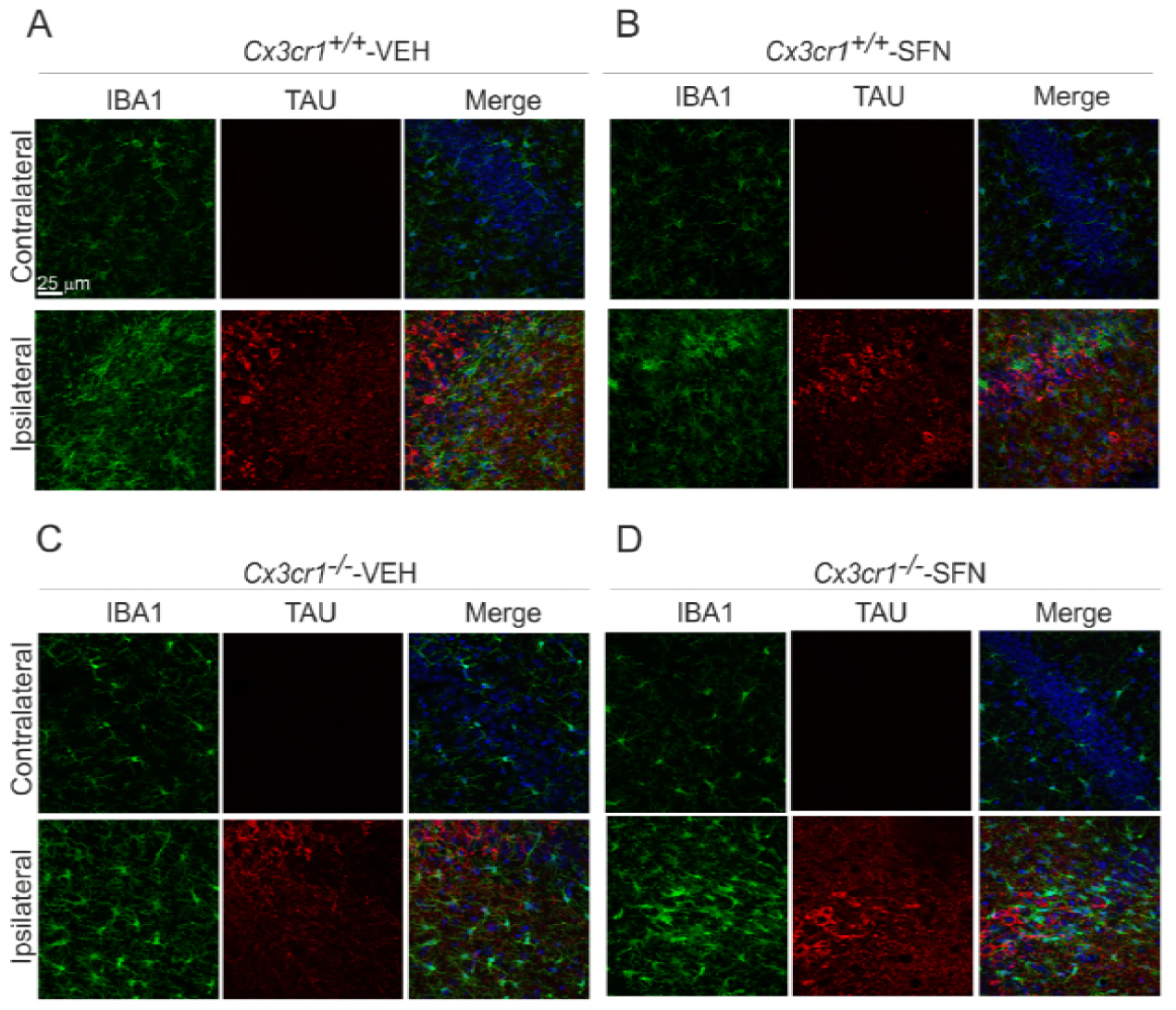
TAUP^301L^ overexpression induced microglial activation of hippocampus from Cx3cr1^+/+^ and Cx3r1^−/−^ mice. Immunohistochemical staining with anti-human-TAU antibody (red) and IBA1 (green) in 30 μm-thick sections of hippocampus of (**A**) *Cx3cr1*^+/+^-VEH, (**B**) *Cx3cr1*^+/+^-SFN, (**C**) *Cx3cr1*^−/−^-VEH and (**D**) *Cx3cr1^−/−^*-SFN.

**Supplementary Figure 4:**
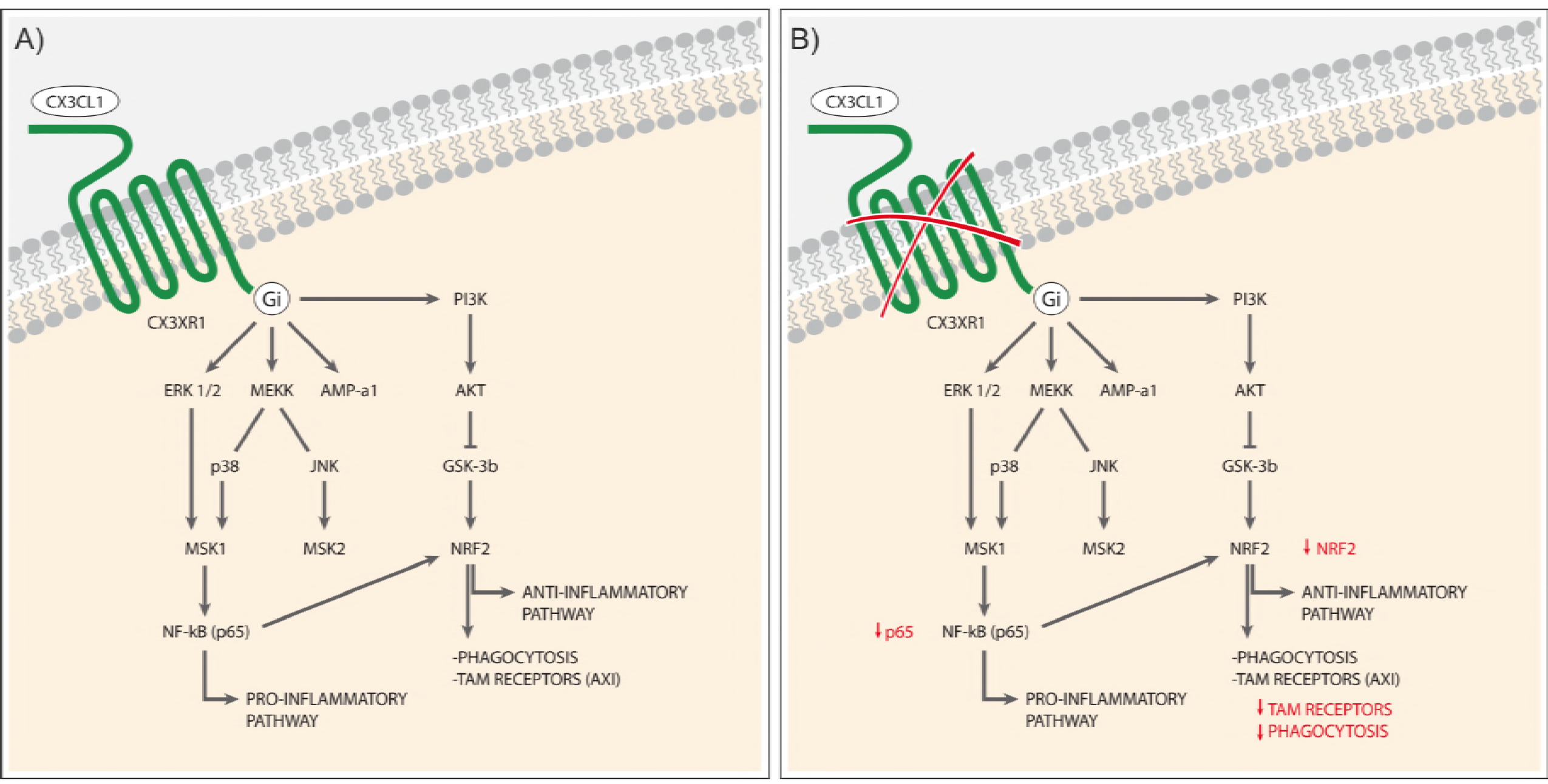
Molecular mechanisms implicated in CX3CL1/CX3CR1 signalling. (**A**) Scheme of the signalling pathways involved in the activation of the transcription factors NF-κB and NRF2, respectively, driven by CX3CL1/CX3CR1 activation. (**B**) Lack of CX3CR1 produces a decrease in the expression of NF-κB and NRF2, and impairment in TAM receptors expression and microglia phagocytosis.

**Supplementary Figure 5:.**
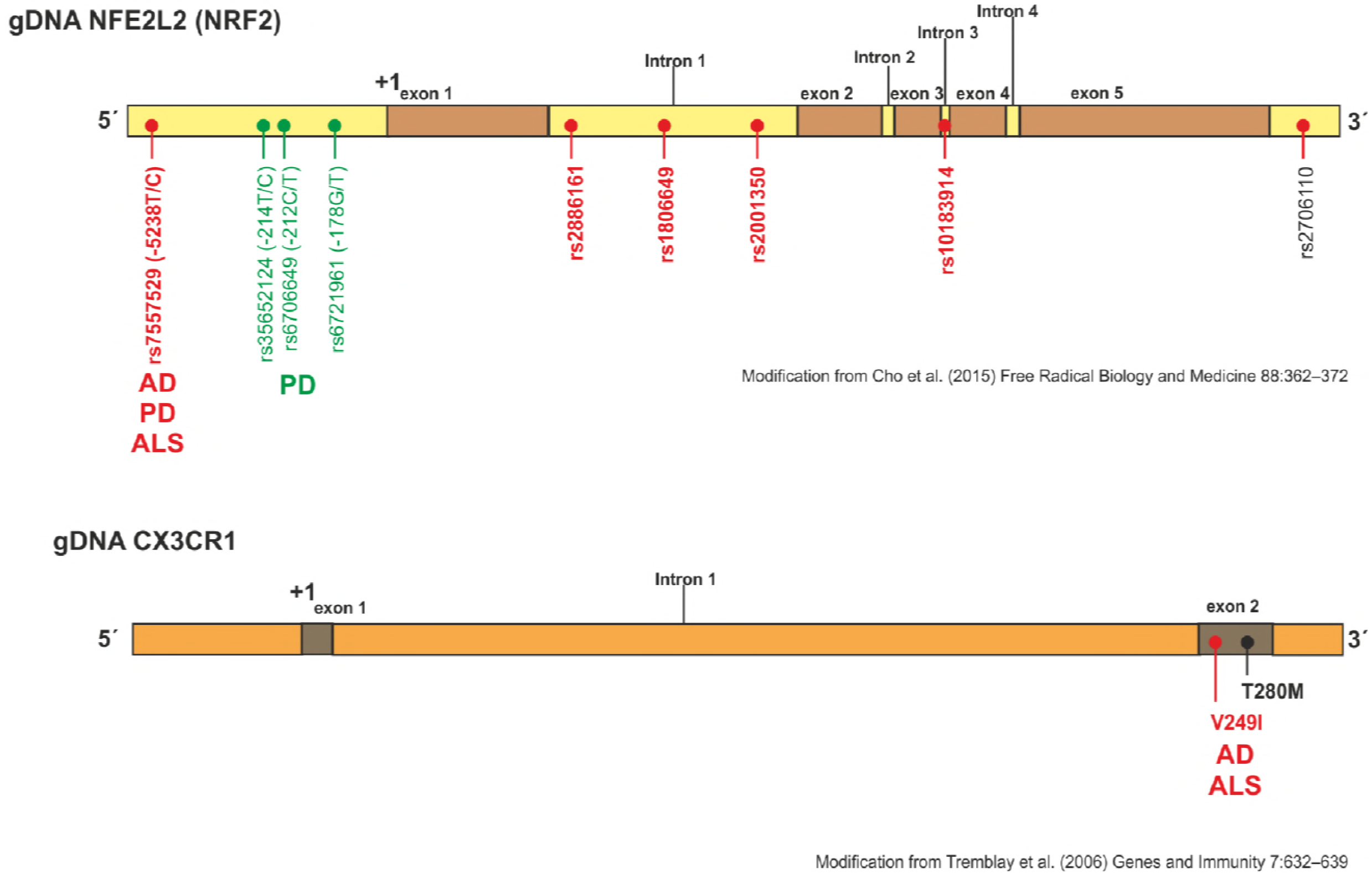
Polymorphisms in NRF2 and CX3CR1 genes associated with neurodegenerative disorders. Scheme of human NRF2 and CX3CR1 genes (modified from Cho et al., 2015 and Tremblay, Lemire et al., 2006 respectively). In this figure, it is depicted the risk of genetic variations in the promoter, exon and introns of the *Nfe2l2* and *Cx3cr1* genes (red dots denote haplotypes that confer increased susceptibility in Alzheimer’s disease (AD), Parkinson’s disease (PD) and Amyloid lateral sclerosis (ALS) whereas green dots show haplotypes that are protective in PD).

